# Australian native flower colours: does nectar reward drive bee pollinator flower preferences?

**DOI:** 10.1101/862110

**Authors:** Mani Shrestha, Jair E Garcia, Martin Burd, Adrian G Dyer

## Abstract

Colour is an important signal that flowering plants use to attract insect pollinators like bees. Previous research in Germany has shown that nectar volume is higher for flower colours that are innately preferred by European bees, suggesting an important link between colour signals, bee preferences and floral rewards. In Australia, flower colour signals have evolved in parallel to the Northern hemisphere to enable easy discrimination and detection by the phylogenetically ancient trichromatic visual system of bees, and native Australian bees also possess similar innate colour preferences to European bees. We measured 59 spectral signatures from flowers present at two preserved native habitats in South Eastern Australia and tested whether there were any significant differences in the frequency of flowers presenting higher nectar rewards depending upon the colour category of the flower signals, as perceived by bees. We also tested if there was a significant correlation between chromatic contrast and the frequency of flowers presenting higher nectar rewards. For the entire sample, and for subsets excluding species in the Asteraceae and Orchidaceae, we found no significant difference among colour categories in the frequency of high nectar reward. This suggests that whilst such relationships between flower colour signals and nectar volume rewards have been observed at a field site in Germany, the effect is likely to be specific at a community level rather than a broad general principle that has resulted in the common signalling of bee flower colours around the world.

## Introduction

Many floral traits play a role in the reproduction of animal-pollinated angiosperms [1–5]. Colour is one of the most important signals used by flowering plants to communicate to their pollinators [6–10]. Flowers typically present nutritional rewards like nectar to entice floral visitors [6,11–15], and nectar is a reward that can promote learning and neural changes in a bees visual system [16]. What relationship between floral colour signals and nectar as a reward that promotes motivation in insect pollinators, should evolve in plants? Whilst nectar has been well studied in flowering plants [15,17–20], the question of a potential relationship has been rarely considered because many animals have very different colour vision to humans [21]. Thus, colour is not an unambiguous trait, and to test colour as a factor in a biologically meaningful way it is necessary to map how relevant pollinators like bees perceive and use colour information.

Both male and female fitness of plants should often benefit from pollinator visits, especially given widespread pollen-limitation of seed set [22]. Such fitness benefits for plants will be especially strong with flower visitors like bees that tend to be ‘flower constant,’ that is, to use colour signals to repeatedly visit flowers of the same plant species [23]. In turn, the ability of pollinators to assess which floral colour signals are more reliable predictors of nutritional rewards will affect the foraging success of individuals [24] and thus the subsequent success of bee colonies [14,25–27]. This leads to an interesting hypothesis that bees may have evolved colour preferences because visiting flowers with higher rewards improves foraging performance while flowers gain increased pollination services by signaling higher rewards in a reciprocal selection loop that promotes the evolution of certain flower colours [28].

Colour vision requires multiple photoreceptors with different sensitivities (Jacobs 2018). The spectral sensitivities of photoreceptors in many bee species have been empirically determined, showing that the trichromatic colour vision of bees is highly conserved and predates the evolution of flowers [29]. To yield colour information, photoreceptor signals have to be antagonistically processed in a brain [21]. Such colour opponent mechanisms in bees have been empirically recorded [30–32]. Knowing this information enables the construction of a colour space that accurately represents colour information, and the colour Hexagon is an opponent colour space that represents the visual capabilities of trichromatic bees [33]. In the current manuscript we use capitals (e.g. BLUE) to convey a region in bee colour space (e.g. see Fig. 1a), and ‘blue’ to refer to how humans typically describe colour stimuli, following the convention proposed by [34]).

**Fig 1:**
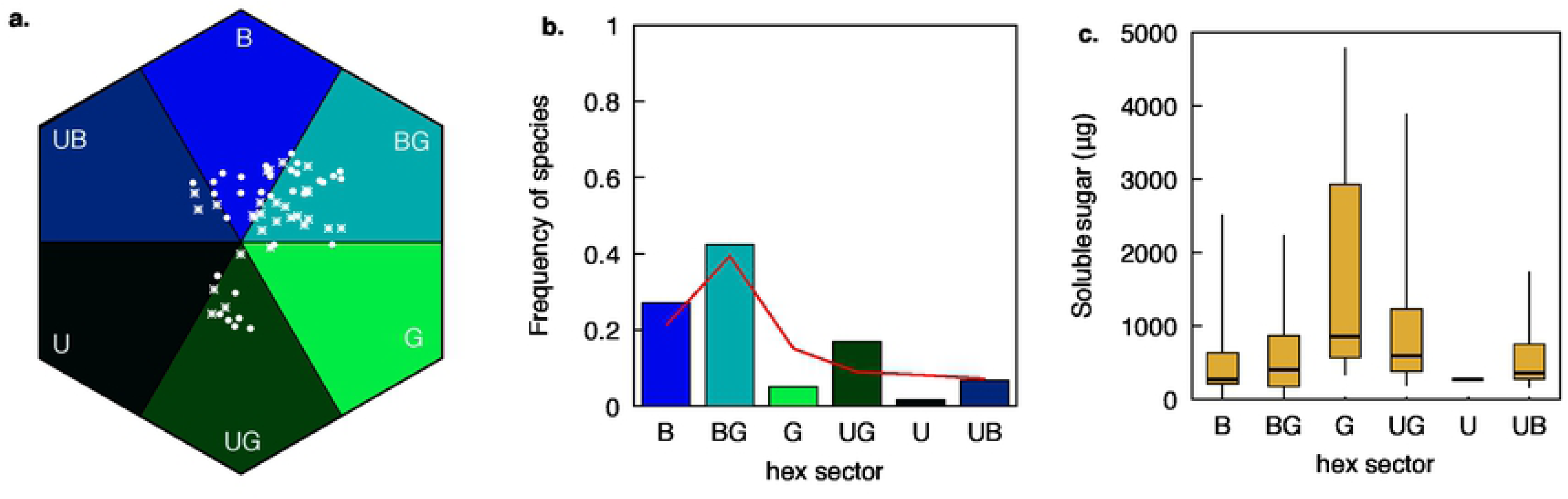
Flower colour and nectar in Australian native plant flowers. **a.** Distribution of 59 flowering plant species in hexagon colour space: non-orchids (•) and orchids (*). b. Frequency of sampled species classified on each of the six Hexagon categories along with the corresponding global pattern of distribution (red line) of plant species taken from the surveys of plant communities in Germany, Australia, and Nepal [34,43,46,56]. c. Plant flower soluble sugars by colour category; thick lines represent medians, boxes represent the 25% and 75% interquartile ranges, and thin vertical bars represent 2.5 and 97.5 % quantiles of the data distribution. Names of the different Hexagon sectors are abbreviated: BLUE (B), BLUE-GREEN (BG). GREEN (G), UV-GREEN (UG), UV (U), and UV-BLUE (UB) as described by [34].

Interestingly, both honeybees and bumblebees show similar distinct preferences for short wavelength ‘blue’ stimuli that frequently have loci in the UV-BLUE, BLUE and/or BLUE-GREEN sectors of bee colour space [28,35–40]. It has also been recently shown that native bees in Australia show a significant colour preference for stimuli in the BLUE and BLUE-GREEN regions of bee colour space [41,42]. These potentially common bee colour preferences provide a plausible explanation for why bee pollinated flowers around the world frequently share similar distributions in colour space [43].

Floral colour distributions in natural plant communities have recently been documented from a geographically wide array of sites [3,44–52] including several recent studies that use sophisticated modeling of pollinator colour vision [43,52–54]. Where bees are present in both the Northern [3,46,55] and Southern [56–58] Hemispheres, flowers tend to have evolved colour signals that are efficiently processed by bee trichromatic vision. In Australia it has been observed at both a continental level [56,58] and a local community level [43,54] that flower colours share a very similar distribution in colour space to flower colour distributions in the Northern Hemisphere where bees with a blue preference are the dominant pollinator. Evolutionary change in floral colour appears not to be strongly limited by phylogenetic constraints [43,46,47,53,58], and it is of high value to understand what does promote very consistent flower colouration around the world when considering the visual capabilities of bees.

In a study at a field site near Berlin, Germany, flowers were most frequently found to be in the UV-BLUE region of colour space and these flowers also contained higher nectar rewards [28]. In the current study, we seek to understand if floral colours at a community level in Australia also show a significant association with nectar rewards that is consistent with the pattern found in Germany [28], and thus whether floral nectar reward might be an important driver linking bee preferences and flower colours.

## Materials and Methods

### Sites and data collection

We collected data from two natural communities in central Victoria, Australia: Baluk Willam Flora Reserve (37°55′32′′S, 145°20′45′′E), 40 km south east of Melbourne, and Boomers Reserve (37°37′39′′S, 145°15′21′′E), approximately 35 km north east of Melbourne. Both sites are *Eucalyptus* woodland with well-developed shrub and herb layers containing a high diversity of orchid species. These communities have been protected to maintain native vegetation. Flowers were sampled from March 2010 to May 2011. Species were identified with the aid of several local floras [59–66]. A list of species included in this study is given in online Appendix A.

### Nectar measurement

We used the floral nectar data of 59 bee pollinated flowering plant species in our analysis. Nectar collection and measurement methods are detailed in [54]. Briefly, newly opened flowers were place in pollinator exclusion nets for 24 hours to allow nectar accumulation. Flowers were then excised, and soluble sugars were extracted by immersing whole flowers in known volume of distilled water followed by an acid treatment to reduce all sugars into hexoses. Subsequently, we measured the concentration of soluble sugars using standard spectrophotometric methods and back calculated the total sugar content of each flower [67]. Quantity of sucrose present in each flowering plant species is listed in online supplementary [54] and Appendix A.

### Colour measurement

Reflectance spectra from 300 to 700 nm wavelength were measured on a minimum of three flowers for each species using an Ocean Optics spectrophotometer (Dunedin, Florida, USA) with a PX-2 pulsed xenon light source. A UV-reflecting white standard was used to calibrate the spectrophotometer. Spectra from multiple flowers were averaged within each species. For flowers with multiple colours, such as areas with and without a UV component, we obtained reflectance spectra of the two colours covering the largest surface area of the flower.

### Colour space representation

Floral reflectance spectra were converted to positions in a hexagonal colour space, a two-dimensional representation of the excitation levels of the three different classes of photoreceptors in a hymenopteran insect’s visual system [33]. This model is widely accepted as a representation of bee trichromatic vision in comparative studies of flower evolution [3,28,34,39,41,43,50,53,55,56,68,69]. The exact photoreceptor sensitivities of native Australian bees are currently unknown, but relying on the phylogenetic conservation of spectral sensitivity peaks of hymenopteran photoreceptors [29], we use a general hymenopteran visual model based on a vitamin A1 template for photopigments [70] with sensitivity peaks at 350 nm (ULTRAVIOLET: UV), 440 nm (BLUE: B) and 540 nm (GREEN: G) (cf. [56]). We calculated the relative probability of photon capture (*P*) by each of the UV, B, and G photoreceptors by numerically integrating the product of the spectral sensitivity function of each one of the (*i* = 3) photoreceptors *S*_*i*_(*λ*), the diffuse spectral reflectance of each flower *I*(*λ*) and the spectral distribution of the ambient illumination *D*(*λ*) expressed as photon flux [71]. All spectral functions were expressed from 300 to 650 nm at 10 nm steps:

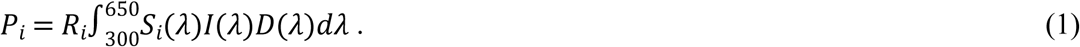

The coefficient *R*_*i*_ in equation 1 represents von Kries adaptation and is used to normalize each of the photoreceptors to the illumination reflected from the background [33]:

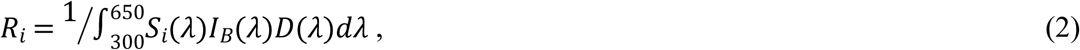

where *I*_B_ (λ) is the spectral reflectance of the background. We used the average reflectance from 20 species of *Eucalyptus* (Average Green Leaf) as background reflectance (*I*_*B*_(λ)) for our calculations. We used a open sky, daylight ambient illumination equivalent to CIE 6,500 K [72], that represent typical daylight conditions for foraging insects [73]. The transduction of photoreceptor captures (*P*) into receptor excitations (*E*) is given by

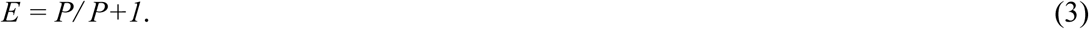

The receptor excitations (*E*_*UV*_, *E*_*B*_ and *E*_*G*_) are plotted on orthogonal axes, each of unit length, and the colour locus of a flower is the vector sum of the individual excitations. A colour locus can be represented by Cartesian coordinates in a hexagonal space using equations 4 [33]:

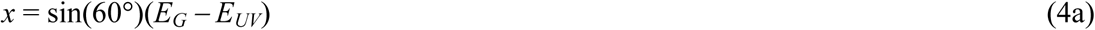

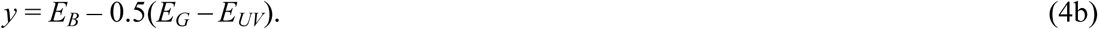

Colour contrast in Hexagon space was calculated as the Euclidean distance from the centre of the colour space, representing the adapting background, to the locus of a flower [74]. Hexagon coordinates for all flower species are given in online Appendix A.

## Data analysis

### Colour categories

For each species, we calculated the polar coordinates (angle and magnitude) of the floral colour locus in the hexagonal colour model (Fig 1a). The angle is a measurement of ‘hue’ in the hexagon space [33]. Samples were subsequently classified into one of the six colour categories proposed by [34] based on their respective hue value: BLUE (B), BLUE-GREEN (BG). GREEN (G), UV-GREEN (UG), UV (U), and UV-BLUE (UB).

### Does sugar content vary among colour categories?

Flowers were subsequently classified as either having a ‘high’ or ‘low’ soluble sugar content relative to the median soluble sugar amount for the entire flower sample, following the method used in [28,75,76]. We used a set of contingency tables to test for significant differences in proportion of high and low soluble sugar content per color category, against a null hypothesis of equality of proportion per color group. Contingency tables excluded the single sample present in the UV color group (see results section for details). To avoid problems associated with using the χ^2^ distribution with small sample sizes in some colour categories, probability values from the chi-square test were obtained using 100,000 Monte Carlo simulations.

A second contingency test was conducted after excluding three species from the family Asteraceae because the soluble sugar content for these species was measured from a compound ‘head’ rather than individual flowers. Median soluble sugar amount was thus recalculated from the remaining species and the response variable was reformulated using the updated soluble sugar threshold value.

Finally, we constructed a third table excluding plant species in both Asteraceae and orchidaceae, given the prevalence of potential food deception in many orchids [77,78]. As before, the median soluble sugar content was updated, and the remaining flower species were subsequently reclassified as being high or low.

### Correlation between sugar content and chromatic contrast

In addition to hue, chromatic contrast with a background (see *Colour space representation* above) is an element of colour that may be relevant to pollinators. We tested for a potential correlation between chromatic contrast of flower loci to the leaf green background in hexagon colour space and soluble sugar content. The analysis was done first on the entire data set, and subsequently for the two data subsets following the same rationale used for the contingency tests. All correlation tests were performed using Kendall’s tau (τ) statistic as the test makes no assumptions on the underlying distribution of the data [79]. All analyses were performed using R base package version 3.6.1 (05-07-2019).

## Results

The distribution of species among hexagon sectors appeared uneven (Fig 1a, b), as did the distribution of soluble floral sugar per sector (Fig 1c). Only one sample was classified into the UV hexagon sector: *Hypericum pygmae* (Clusiaceae) and this hexagon sector was thus excluded from all subsequent analyses. The scarcity of flowers in UV sector is consistent with previous studies [34].

Median soluble sugar content per flower ± median absolute deviation (MAD) for the sample excluding *Hypericum pygmae* was 392 ± 377 μg. Following categorization of species based on this threshold value (Fig 2a), we found no significant difference among hexagon sectors in the proportion of species with a high sugar content (χ^2^ = 3.97, *P* = 0.466).

**Fig 2:**
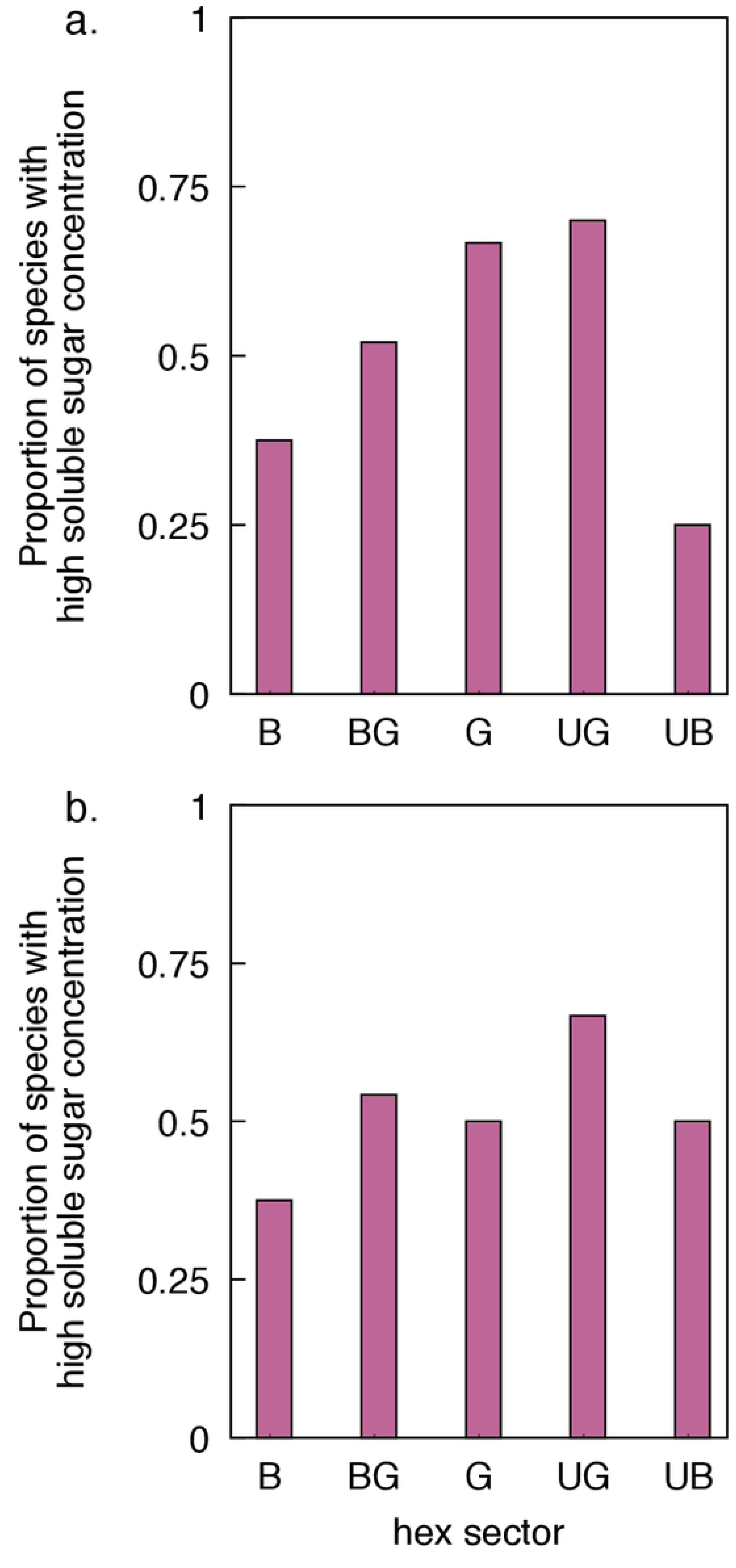
Proportion of species with a ‘high’ amount of soluble sugar for each one of the five categories including (panel a) and excluding species from the family Asteraceae (panel b).

Results from the second contingency table (median sugar soluble sugar = 383 ± 359 μg), which excluded species from the family Asteraceae also failed to reject the null hypothesis of equality in the proportion of species with a high amount of soluble sugar (χ^2^ = 2.15, *P* = 0.765, Fig 2b).

Finally, the threshold soluble sugar value for non-orchid, non-aster species contingency table was 367 ± 406 μg. For the third model we also found no evidence rejecting the null hypothesis of equality in the proportion of high-reward species among the different colour categories (χ^2^ = 2.97, *P* = 0.693). This result thus suggests that the low amount of soluble sugars present in orchid species at our field site was not a factor affecting a potentially the outcome of the initial model.

Chromatic contrast revealed no significant relation to floral sugar content at our field site. Tests using Kendall’s tau (τ) statistic failed to reject the null hypothesis of independence between soluble nectar content and chromatic contrast for all the subsets considered (Table 1).

**Table 1:** Results of correlation test testing for a potential relation between soluble sugar content and chromatic contrast considering various sample subsets: Complete data set including all flowers, data set excluding the only species allocated to the UV hexagon sector *Hypericum pygmae*, subset also excluding family Asteraceae, and all non-orchid species. A non-parametric (Kendall tau (τ)) correlation coefficient was calculated in all cases.

| Data set | $\tau$ | P |
| --- | --- | --- |
| Complete data set | -0.016 | 0.86 |
| Excluding flower in UV sector | -0.019 | 0.83 |
| Excluding Asteraceae | -0.059 | 0.528 |
| Excluding Orchidaceae | 0.003 | 0.999 |

## Discussion

It was hypothesised by Darwin [80] that insects may evolve innate preferences to aid the efficient location of profitable flowers, and bees do have both innate spatial [5,81] and colour [28,35–40] preferences. In Germany, where important bee species like honeybees and bumbles have innate preferences for short wavelength blue flowers, it was found that flowers in the UV-BLUE category contained significantly higher volumes of nectar than those in other hue categories in bee colour space [28]. Several studies suggest that nectar volume can influence the behaviour of foraging bees [11,12,19,20,82]. Indeed, it has also been shown that introduced species with flowers that contain higher volume rewards can out-compete resident flowers by attracting more bee pollinators [83].

Nonetheless, flower colours in different parts of the world have very similar distributions in bee colour space [56], which is also consistent with evidence that bees have phylogenetically conserved colour visual systems including preferences for short wavelength stimuli [29,43]. Thus, understanding whether flowers in different communities have colours that predict higher reward levels in a consistent fashion is of value for understanding what traits promote bee choices, and the potential major drivers that influence flower signal evolution.

In the current study we considered flower colour signaling from two communities in south-eastern Australia that had similar flower colours in bee colour space, and were also similar to bee pollinated flower colours found elsewhere across Australia and around the world [3,43,46,55,56]. We found no evidence that a particular hue predicted a higher reward level within flowers. These results from native plant communities suggest that a simple, direct link between flower rewards and the preferred bee hue of ‘blue’ is not an explanation of flower colour preferences in Australia, and may not be in other locations around the world. Reciprocal selection between plants and their pollinators can reach different evolutionary equilibria among local populations, leading to a geographic mosaic of trait values of the interacting parties [84]. The previously observed higher rewards for bee preferred blue flowers in Germany [28] may thus be a local equilibrium, and it will be of value to map more communities to understand if and how flower colour predicts nectar rewards around the world.

Given the evidence that bee colour preferences may influence how flowers evolve similar spectral signals at several different locations around the world [34,43,46,56,57,85], it is interesting to consider what traits other than nectar rewards might promote bee preferences. Plausible alternative lines of investigation could include how the spectral overlap of photoreceptors when combined with opponent processes at a neural level enhance both colour discrimination [55,86–88] and colour detection [89] in a way that is most efficient for finding flowers [90]. This in turn could enhance neural mechanisms to promote innate colour preferences. By itself, this mechanism of spectral tuning cannot be the sole explanation for a stronger blue preference, since there is also spectral overlap and enhanced signal processing at longer wavelengths [55,85]. However, many common background stimuli reflect at longer wavelengths [91], so having innate brain preferences for shorter wavelength ‘blue’ stimuli might enable bees to efficiently detect stimuli that have a very high probability of being a rewarding flower given that very few natural colours are blue. Interestingly, UV absorbing flowers that appear ‘white’ to humans are very common within this short wavelength range of preferred colours in bee colour space, and additionally have the advantage of having strong modulation of the long wavelength bee receptor that is implicated in enhancing signal detection at a distance [68,92–95].

To better understand the complexities of these multiple complex factors it is important to collect more flower data at a plant community level around the world, as well as continue to map the sensory capabilities of different pollinators [16,87].

## Acknowledgements

We thank Parks Victoria and the Department of Sustainability and Environment (permit number 10005294) for permission to work at the field sites between 2009-2013. We thank Professor Graham H. Pyke for the suggestions and comments about nectar rewards in the previous version of manuscript.

## Author contributions

MS, AGD and MB designed the study. MS conducted the field work, data collections, plant identification. MS and MB did the data curations and lab experiment. MS, and AGD mapped the floral reflectance spectra to the bee-vision colour space model. MS, JEG, MB, AGD performed the statistical analyses. All authors interpreted the results and wrote the manuscript. The authors declare that they have no conflict of interest involving the work reported here.

## Supplementary information

## Appendix A

provides the data used in current analysis. The ‘.csv’ file includes hexagon x and y unit, sucrose amount (microgram) and pollination categorization.

